# Laser ablation of the apical sensory organ of *Hydroides elegans* (Polychaeta) does not inhibit detection of metamorphic cues

**DOI:** 10.1101/2021.01.23.427931

**Authors:** Brian T. Nedved, Marnie L. Freckelton, Michael G. Hadfield

**Affiliations:** University of Hawaii, Kewalo Marine Laboratory, Honolulu, 96813, United States

**Keywords:** apical sensory organ, *Hydroides elegans*, larval development, metamorphosis

## Abstract

Larvae of many marine invertebrates bear an anteriorly positioned apical sensory organ (ASO) presumed to be the receptor for settlement- and metamorphosis-inducing environmental cues, based on its structure, position and observed larval behavior. Larvae of the polychaete *Hydroides elegans* are induced to settle by bacterial biofilms, which they explore with their ASO and surrounding anteroventral surfaces. A micro-laser was utilized to destroy the ASO and other anterior ciliary structures in competent larvae of *H. elegans*. After ablation, larvae were challenged with bacterial biofilmed or clean surfaces and percent metamorphosis was determined. Ablated larvae were also assessed for cellular damage by applying fluorescently tagged FMRF-amide antibodies and observing the larvae by laser-scanning confocal microscopy. While the laser pulses caused extensive damage to the ASO and surrounding cells, they did not inhibit metamorphosis. We conclude that the ASO is not a required receptor site for cues that induce metamorphosis.

**Summary Statement:** Larvae of the polychaete *Hydroides elegans* retain the capacity to sense biofilm cues and metamorphose despite removal of their apical sensory organs, the supposed sensors for settlement cues.

## Introduction

The communities of animals that inhabit the benthos in the seas, from the highest reaches of the seashore to the greatest depths, depend on the successful recruitment of very small larvae for their establishment and maintenance. A large majority of these animals produce larvae that spend hours to months swimming and growing in the plankton before they are developmentally competent to settle and metamorphose. At this point, they must locate an appropriate site for settlement, one that will provide the right habitat requirements be it mud, sand, or rocks, and access to appropriate food and other individuals of their species with which to mate and produce new generations. Each of these very small, ready-to-settle larvae, equipped with few simple sensory organs, must detect unique cues from habitats and settle selectively onto only the ‘right one.’ Critical questions regarding site-specific larval recruitment concern how and where, on their bodies, larvae detect habitat-specific environmental cues.

Major focus for a wide variety of invertebrate larvae has been on the apical sensory organ (ASO), a feature of many different types of larvae across many phyla (Nielsen 2004; Hadfield, 2011; Marlow et al., 2014). The ASO is a principal cellular structure of these free-swimming larvae (Wanninger, 2009; Marlow et al., 2014). Its development is typically complete at the onset of larval swimming, either as they develop in the plankton or are released from parental broods or benthic egg masses, and it degenerates at the onset or quickly after the metamorphosis (Marlow et al., 2014; Nelson et al., 2017; Hartingzen, 2018). ASOs are considered homologous across lophotrochozoan phyla, and representatives of unrelated taxa within the cnidarians and bilaterians have ASOs of similar structure (Chia and Koss, 1979; Nielsen, 2004; Byrne et al., 2007; Nielsen 2008; Miyamoto et al., 2010; Rawlinson 2010; Hadfield, 2011; Hindinger et al., 2013; Marlow et al., 2014). Even some highly modified larvae, which do not swim and undergo metamorphosis inside their egg capsules, have this organ (Voronezhskaya and Khabarova, 2003). The ASO is always located at the anterior pole of the larva that is directed forward during swimming. Its characteristic features are a tuft of sensory cilia arising from a cluster of sensory neurons that lie just anterior to major ganglia (Conklin, 1897; Beklemishev, 1964; Page, 2002; Yuan et al., 2008; Marlow et al., 2014). The cells from which the tuft arises are serotonergic or FMRF-amidergic in most lophotrochozoans where it has been investigated (reviewed by Lacalli, 1994 and Marlow et al., 2014). The unique evolutionary constancy of the ASO indicates that its function may be conserved and shared across many larval types.

For a few invertebrate species, there is evidence that the ASO is the receptor for a specific stimulatory ligand (e.g., Hadfield et al., 2000). As they prepare to settle to the benthos, larvae of many phyla behave in a manner that suggests they are testing or sampling the substratum (Segrove, 1941; Marsden and Anderson, 1981; Gabilondo et al., 2013; Nelson et al., 2017). Typically, this pre-settlement behavior includes swimming close to the surface at slower speeds than when swimming in the plankton and orienting themselves with the apical end toward the substratum and the apical tuft brushing the surface (Barnes and Gonor, 1973; Nott, 1973; Marsden and Anderson, 1981; Zimmer, 1991; Gabilondo et al., 2013; Nelson et al., 2017). Thus, the apical tuft and its associated cells have long been suspected to be the site of detection of cues for settlement and metamorphosis (Hadfield, 2011).

The circum-globally distributed, warm-water serpulid polychaete *Hydroides elegans* (Haswell) has emerged as a useful model for studying larval settlement phenomena, especially induction by specific biofilm bacterial species (Hadfield et al., 1994; Unabia and Hadfield, 1999; Lau and Qian, 2002; Huang and Hadfield, 2003; Nedved and Hadfield, 2009; Shikuma et al., 2014; Freckelton et al., 2017; Vijayan and Hadfield, 2020). Evidence for detection of a cue is strong: larvae of *H. elegans* typically will not settle in the absence of a biofilm, and induction of metamorphosis by bacteria requires contact with a biofilm (Hadfield et al., 2014). Further, while numerous biofilm bacterial species will induce metamorphosis in larvae of *H. elegans*, some – perhaps most – biofilm-bacterial species do not (Unabia and Hadfield, 1999; Lau and Qian, 2001; Vijayan and Hadfield, 2020).

There are at least three potential sites for chemoreception of bacterial cues on the episphere of competent larvae of *H. elegans*. The first site, the apical sensory organ (ASO), is prominent at the anterior apex of the episphere. An apical tuft of sensory cilia (AT) projects from the ASO into the environment (Figure 1A). There are four sensory cells in the ASO and an additional 12-14 cells in the cerebral ganglia that are strongly immunoreactive to polyclonal antibodies raised against the neuropeptide FRMF-amide (Nedved, 2010). Lying to each side of the ASO are two additional sensory patches. Most distal are the lateral sensory organs (Fig. 1A, LSOs), each including two, cilia-bearing, FMRF-amide positive cells (Fig. 1B). The medial sensory organs (Fig. 1B, MSOs) lie between the LSOs and the ASO. One sensory cell within the MSO is FMRF-amide positive and projects a fiber into the cerebral ganglia.

**Figure 1.**
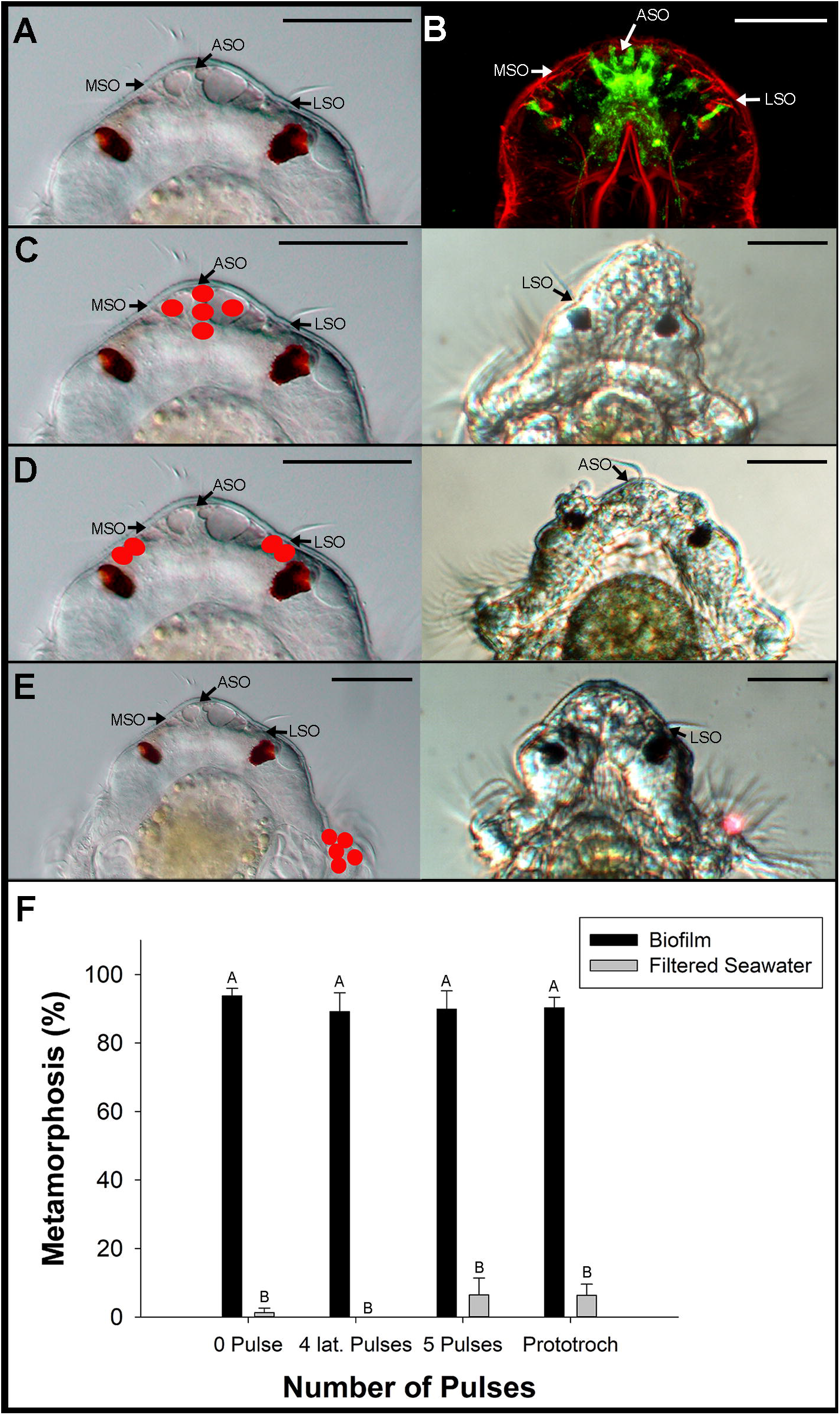
Location of targeted cells and damage caused by laser ablations of the anterior sensory organ (ASO) and surrounding sensory cells in larvae of *Hydroides elegans*. (A) Morphology of the episphere of a competent larva. (B) Location and morphology of FMRFa-immunoreactive cells (green) in the episphere of a competent larva. (C) Left, locations of five laser pulses administered to ASO; right, morphological damage caused by 5 pulses. (D) Left, location of 4 laser pulses administered to LSOs; right, damage caused by 4 pulses to LSOs. (E) Left, location of 5 laser pulses administered to the right side of the prototroch; right, damage caused by five pulses to prototroch. (F) Percentage of larvae that metamorphosed 14-16 h after ablation of the cells within the ASO (5 pulses), LSOs (2 pulses on each), and cells in the prototroch (5 pulses). N=4 replicates. The experiment was performed once. Kruskal Wallace test, p-value <0.0006. A Dunnett’s test was used to compare treatments against the control treatment (Biofilm, 0 Pulse). Bars=means, Error bars =1 SE. Bars with the same letter are not significantly different from the control treatment. ASO= apical sensory organ, MSO= medial sensory organ, LSO = lateral sensory organ. Scale bars = 25 µm.

It was the goal of the research presented here to experimentally test the hypothesis that the ASO is the receptor for bacterial settlement cues for larvae of *H. elegans*. The approach employed micro-laser destruction of the targeted ciliary organs (ASO, MSOs, and LSOs) followed by assays using FMRF-a antibodies to determine the extent of the damage, and settlement assays to determine if larvae with destroyed sensory organs could still detect bacterial sensory cues and respond to them with settlement and metamorphose.

## Materials and methods

### Culture of larvae

Adults of *Hydroides elegans* were collected from Pearl Harbor, HI (21°21’25.5” N, 157°57’35.9” W) and maintained in flowing seawater at the Kewalo Marine Laboratory of the University of Hawai□i at Mānoa. Worms were induced to spawn by removal from their tubes. Fertilization occurred in 0.22 μm filtered seawater (FSW), and development progressed to the feeding trochophore stage in about 12 h. Larvae were cultured using the methods of Nedved and Hadfield (2009). Briefly, larvae were maintained in FSW at a density of 10 larvae ml^-1^ at 25-26°C. Larvae were fed the single-celled alga *Isochrysis galbana* (Tahitian strain) at a concentration of 6 × 10^4^ cells.ml^-1^. The culture vessels and FSW were changed daily to minimize buildup of bacterial films. Larvae became metamorphically competent 5 days post-fertilization; competent larvae were used in all experiments and assays.

### Laser ablation of sensory cells

Larvae were relaxed in 3.7% MgCl_2_ in FSW. Five larvae were pipetted in a small volume of seawater onto a Rain-X^®^ coated microscope slide. Larvae were covered with a #1 coverslip that had its edges supported by a double thickness of Scotch^®^ tape, which served to immobilize the larvae. By moving the coverslip, larvae were rolled until their dorsal sides were facing up. Cell ablations were performed using a XYclone infrared laser (Hamilton Thorne, Beverly, MA) mounted to a 20X objective on a Zeiss Axiophot upright microscope at 100% power. The laser was aimed at the selected location on the larva and 500 μs pulses were fired the desired number of times (Fig. 1). This procedure was then repeated for each larva on the slide. Two different types of controls were utilized in all ablation experiments. A laser control was implemented by focusing on the right side of the prototroch and administering five pulses. A set of handling controls was also implemented. For these controls, a subset of larvae was pipetted, immobilized and rolled on a slide. No laser pulses were applied in this treatment. After ablation, the coverslips were carefully removed, and the water droplet containing the ablated larvae was removed from the slide by pipetting. Larvae were deposited into dishes of FSW and allowed to recover for 1 h before use.

### Induction of metamorphosis in treated and untreated larvae

After ablation, larvae were split into two treatments: (1) exposed to a complex natural bacterial biofilm at least two weeks old or, (2) kept in FSW, both in 35 mm petri dishes. Four replicate dishes were used for each treatment. The proportion of larvae that had metamorphosed was determined after 14-16 h. Proportional data were transformed by the arc-sine of the square root of the variate (Sokal and Rohlf, 1981). Transformed data failed the Shapiro-Wilks test for normality, and the transformed means were compared using the non-parametric Dunnet’s test using the Analyse-it (Ver. 5.65) statistical software for Microsoft Excel (Analyse-it Software, Ltd. https://analyse-it.com). Means and standard errors were back-transformed to percentages and plotted in SigmaPlot (Ver 12.3, Jandel Scientific Software Inc.).

### Visualizing damage to sensory cells

The immunocytological methods described below were modified from Croll and Chiasson (1989) to determine the expression of the neurotransmitter FMRF-amide in the developing larvae of *H. elegans*. FMRF-amide antibodies were chosen because they label cells of the ASO and as well as sensory cells in the LSOs and MSOs, plus cells within the cerebral ganglia, and axons in the cerebral commissure.

The laser-ablation methods described above were administered to additional larvae. After recovering, each larva was pipetted into a separate well of a 24-well plate that also contained a chip of biofilmed microscope slide. Larvae were monitored once an hour, and if a larva had begun to metamorphose and had secreted a primary tube, the chip was transferred to a dish containing ice-cold 3.7% MgCl_2_ in FSW for 5 minutes to anesthetize the larva. Although these larvae had irreversibly begun the metamorphic cascade, they still retained the ASO, MSO and LSO sensory complexes. After larvae were relaxed, an eyelash brush was used to gently coax each metamorphosing larva from its primary tube. Larvae were then fixed for 1 h in 4% paraformaldehyde in FSW (4°C). After fixation, larvae were washed (3 × 5 min) in phosphate buffered saline (PBS; 50 mM Na_2_PO_4_, 140 mM NaCl pH 7.4). Larvae were then washed (3 × 30 min) in PBT (PBS + 0.01% Triton X100) and further incubated for 20-30 min in blocking solution (PBT + 5% v/v heat inactivated normal goat serum). Larvae were incubated for two days at 4°C in the polyclonal antibody anti-FMRF-amide (raised in rabbit, DiaSorin, Stillwater, MN), at a dilution of 1:500 (v/v in blocking solution). After incubation in primary antibodies, larvae were subjected to three, 2-3 min washes with PBT, followed by three washes (30 min) in PBT, and then incubated in blocking solution (30 min). Larvae were subsequently incubated for 12-24 h at 4°C in Alexa Fluor^®^488 goat anti-rabbit (1:1000 dilution, Molecular Probes). After incubation in the secondary antibodies, larvae were washed several times in PBS, immersed overnight in 3:1 glycerol:PBS, and mounted on glass microscope slides in the PBS:glycerol solution.

In order to visualize the position of labeled nerves within each larva, fluorescently tagged phalloidin was used as a counterstain to provide morphological landmarks within the larvae. A 1:500 dilution of Alexa Fluor^®^ 594 phalloidin (Molecular Probes) was co-incubated with the secondary antibody solutions. Labeled larvae were examined using a Zeiss LSM 710 scanning confocal microscope (Zeiss, Germany) equipped with the appropriate laser and filter combinations. Digital images of optical sections of the preparations were produced using the Zen imaging software package (Version 4.3, Zeiss) and single plane projections of these images, which have a greater depth of focus than single images, were produced using the same software package. Composite plates of these images were constructed using Adobe Photoshop CC2019 (Adobe Systems, San Jose, CA), and the brightness and contrast of each figure was adjusted to provide consistency within the plate.

## Results and Discussion

### Ablation of the ASO and MSOs

Five laser pulses administered to the ASO region caused extensive damage to it, the apical tuft, and the MSO. At most, only one FMRF-a positive cell remained in metamorphosing worms (Figs. 1C,2C and 2F). Despite loss of the cells underlying the apical-tuft and MSO cilia, 91% of ablated larvae completed metamorphosis when exposed to a natural biofilm (Fig. 1F).

### Ablation of the LSO

Four laser pulses administered to the LSOs ablated all the lateral sensory cells in 73% of larvae examined (Fig. 1D and 2B). In eight larvae, at least one (of two on each side) FMRF-a positive cell associated with either the left or right LSO. These lateral pulses left the ASO, MSO, and the cerebral ganglion undamaged but destroyed both the LSO and its sensory cilia in all preparations. (Figs 2B and 2E).

**Figure 2.**
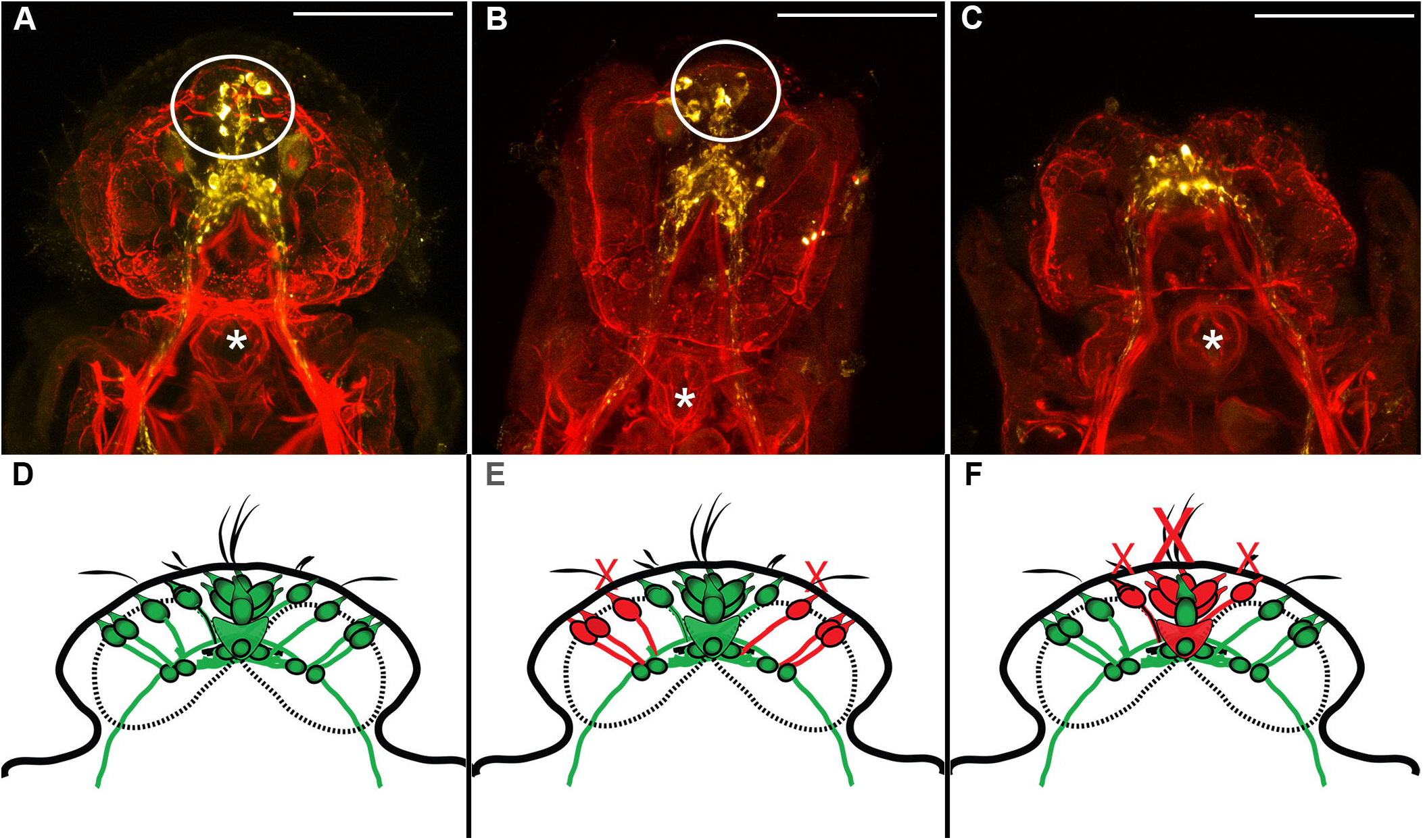
Cellular damage by laser ablation in the apical sensory organ, medial sensory ciliated organs, and lateral sensory ciliated organs in *Hydroides elegans*. Yellow cells are FMRF-a immunoreactive. (A) Anti-FMRFamide immunoreactive cells (anti-FMRFa) in a metamorphosing larva after five laser pulses to prototroch. (B) Anti-FRMFa cells after 4 lateral pulses damaged sensory cells underlying lateral sensory cilia and medial sensory cilia. (C) Anti-FRMFa cells after 5 laser pulses caused extensive damage to the ASO. Preparations were co-labeled with phalloidin (red). White circles indicate the position of ASO. Asterisks show the position of the larval mouth. (D) Schematic diagram of anti-FMRFa cells in an intact larva. N=30 (E) Schematic diagram of damage caused by 4 lateral laser pulses. N=30 (F) Schematic diagram of damage caused by 5 laser pulses to the ASO and MSOs. N=36 Green cell bodies and fibers are anti-FMRFa positive and undamaged. Red-colored cells were destroyed by laser pulses. Red Xs overlay cilia destroyed by laser pulses. Scale bars = 50 µm.

A large percentage of larvae (90%) that had been exposed to the lateral laser pulses completed metamorphosis when exposed to a natural biofilm (Fig. 1F).

### Ablation of Prototroch Cells

The control laser ablations for these experiments targeted cells in the prototroch, which are neither sensory nor FRMF-amide immunoreactive. The episphere was not damaged (Fig 1E), and 93% of these larvae underwent metamorphosis when exposed to a natural biofilm, a result that is consistent with the response of larvae that have not been subjected to a laser pulse (Fig. 1F).

### Ablation of all Known Sensory Cells in the Larval Episphere

Because ablation of neither the LSOs, MSOs nor the ASO inhibited sensing of metamorphic cues and induction of metamorphosis, we used nine laser pulses to ablate all five of these sensory organs in individual larvae (Fig. 3A and 3B). Survival of these larvae was high and most significantly, nearly all larvae pulsed with the laser metamorphosed when exposed to a complex marine biofilm (Fig. 3C). That is, destruction of the ciliary organs hypothesized to be the sites of bacterial cue detection was not effective in reducing the larval response to the cue. It is also clear from the results that the laser treatments and the loss of neurons within the ASO did not, alone, induce settlement and metamorphosis.

**Figure 3.**
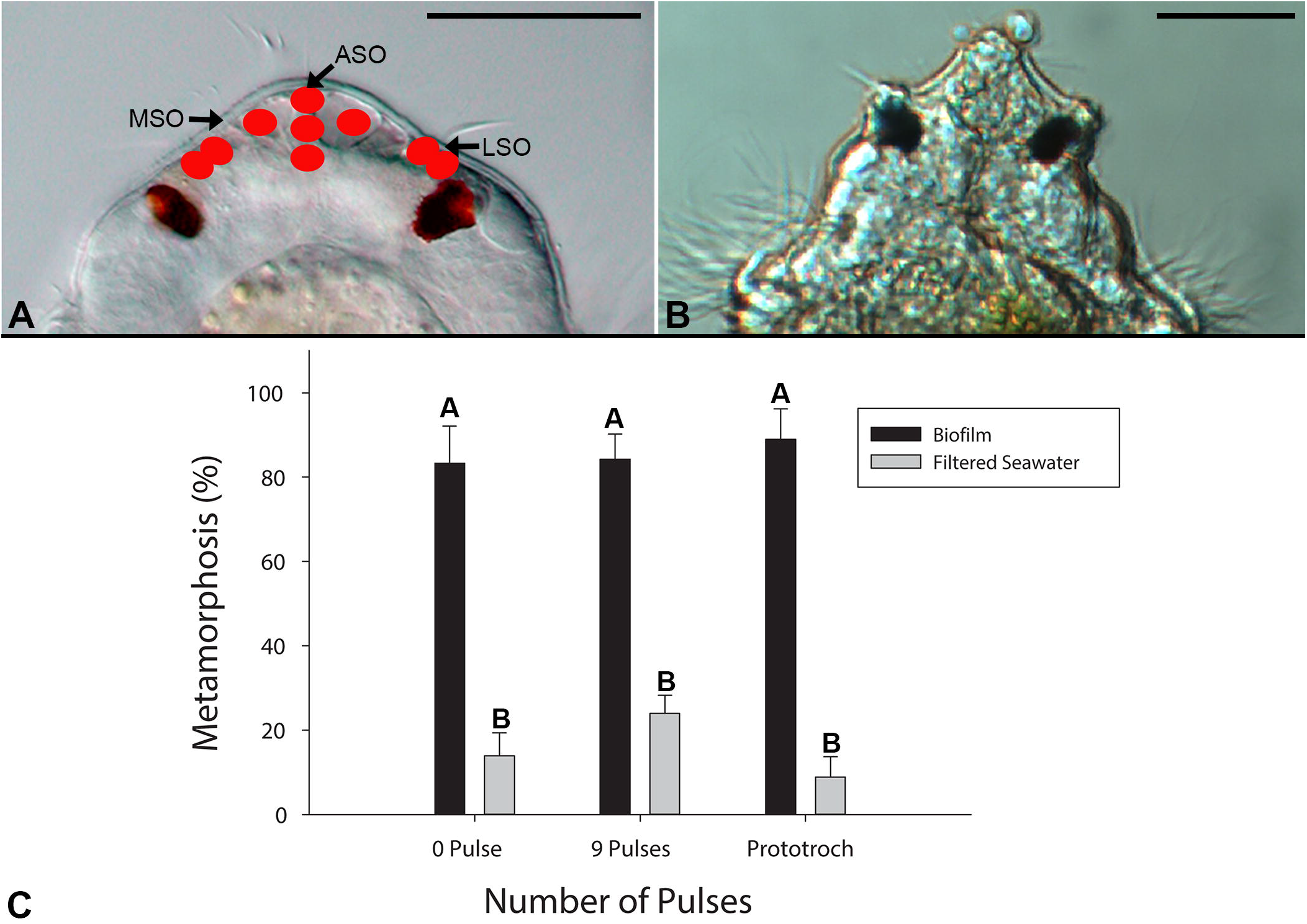
Destruction of the ASO and surrounding tissues does not prevent larvae of *H. elegans* from metamorphosing. (A) Location of nine laser pulses administered to the episphere of larvae. (B) Damage caused by nine laser pulses. (C) Percentage of larvae that metamorphosed after laser ablation. 0 pulses = handling controls; 9 pulses = larvae receiving 9 laser pulses to episphere. Prototroch = larvae receiving 9 laser pulses to right side of prototroch. N=4 replicates. The experiment was performed once. Kruskal Wallace test, p-value =0.002. A Dunnett’s test was used to compare treatments against the control treatment (Biofilm, 0 Pulse). Bars=means, Error bars =1 SE. Bars with the same letter are not significantly different from the control treatment. ASO= apical sensory organ, MSO= medial sensory organ, LSO = lateral sensory organ. Scale bars = 25 µm.

Because laser ablations of the anterior sensory organ (ASO) and nearby ciliated sensory cells did not prevent larvae of *Hydroides elegans* from sensing metamorphic cues we conclude: (1) if the ASO is a detector for bacterial settlement cues, it is not alone in this respect; other receptors must be present in other anterior regions of the larval body; and (2) the function of the ASO may be for a different sensory modality, discussed below. Given that pre-settling larvae of *H. elegans* search for cues by swimming across substrata with the anteroventral regions of their bodies pressed against the surface, it may be that bacterial-cue receptors are located on cilia of the ventral portions of the prototroch, metatroch or food groove, especially around the mouth. We have observed that bacteria are actively swept into the mouth by the actions of prototroch, metatroch and food-groove cilia (Vijayan et al. 2020), and thus it is possible that settlement-cue receptors lie inside the mouth. This location has previously been suggested by Biggers et al (2012) for the polychaete *Capitella telata*. However, all of these regions would be difficult to ablate without other major damage. Also, it remains possible that the anterior sensory structures deleted in our experiments do detect bacterial cues, but not exclusively. Perhaps the larvae of *H. elegans* have multiple receptors located around their pretrochal and ventral postrochal regions.

Based on both developmental and molecular evidence, the ASOs of annelid and molluscan larvae are considered homologous structures (Nielsen, 2004, Marlowe et al., 2014). The ASO of veliger larvae of the nudibranch *Phestilla sibogae* is the sensor for metamorphic cue, and its ablation prevents larvae from sensing the cue and undergoing metamorphosis (Hadfield et al., 2000). However, this function for the ASO may not be completely conserved even within the gastropods. Based on observations made on pharmacologically manipulated, encapsulated, larval-stage embryos of freshwater snails, Voronezhskaya and Khabarova (2003) proposed that the main function of the ASO was to inhibit metamorphosis until an appropriate moment. A similar function for the ASO was proposed for larvae of *Tritia (Ilyanassa) obsoleta*, i.e., a subset of cells within the ASO acts as an inhibitor of programmed-cell death of the ASO, and stimulation of these cells by a mixture of cues from bacteria and diatoms triggers both metamorphosis and the destruction of the ASO (Frogget et al., 1999; Leise et al., 2004; Hens et al., 2006). If the ASO of *H. elegans* serves a similar function, its removal should result in metamorphosis, which we did not observe in our experiments (Fig. 3C).

If the ASO is not a receptor for settlement cues, what is its role in the biology of the larvae of *H. elegans*? Marlow et al. (2014) proposed that ASOs developed as multimodal structures that may have been coopted for different functions in different lineages. Our observations of the apical cilia of swimming larvae of *H*. revealed that the apical cilia are stiff and that they bend when a larva makes a turn (unpublished). It is thus possible that the organ functions in positional sensing in a turbulent world (see Koehl and Cooper, 2015), rather than, or in addition to, chemo-sensing. That is, the ASO may be a mechanical sensor that provides the larva with positional information as well as detecting contact with a solid surface. Marsden and Anderson (1981) also suggested that the ASO may act as a mechanoreceptor in larvae of the serpulid polychaete *Galeolaria caespitosa* where they observed that contact of the apical tuft causes larvae to stop swimming and change direction. Neuropeptides produced by sensory cells within the ASO of larvae of *Platynereis dumerillii* directly influence the speed of ciliary beat in its prototroch and may directly influence the position of the larva in the water column (Conzellmann et al 2011). Indeed, it is possible that the ASO in *H. elegans* is important in mechano- and chemo-sensing of settlement sites, but impulses from the ASO alone do not provide the ultimate signal to attach and metamorphose, a role for additional, yet undiscovered anteroventral receptors.

## Acknowledgements

The authors thank Drs. Audrey Asahina and Elizabeth A. Perotti for their assistance with the ablation experiments.

## Author Contributions

Conceptualization: B.T.N, M.G.H; Methodology: B.T.N., M.G.H.; Investigation: B.T.N.; Analysis: B.T.N., M.L.F.; Data curation: M.L.F., B.T.N.; Writing: B.T.N., M.L.F., M.G.H.; Writing - review & editing: B.T.N., M.L.F., M.G.H.; Supervision and Resources: M.G.H.; Funding acquisition: M.G.H, B.T.N.

## Data Availability

Data are available from FigShare: DOI: 10.6084/m9.figshare.13585052 10.6084/m9.figshare.13585172

